# Synthetic cell armor made of DNA origami

**DOI:** 10.1101/2023.02.20.529284

**Authors:** Weitao Wang, Peter R. Hayes, Xi Ren, Rebecca E. Taylor

## Abstract

Therapeutic and bioengineering applications of cells, such as cell printing and cell delivery, are directly limited by cell damage and death due to harsh mechanical conditions. Improved cellular robustness thus motivates investigations into cell encapsulation that provides essential protection. Here we target the cell-surface glycocalyx and crosslink two layers of DNA origami nanorods on the cellular plasma membrane to form a nanoscale protective shell. This modular and programmable approach enables fine control over the layering and composition of membrane-deposited nanorods. We show that the DNA origami nanoshell modulates the biophysical properties of cell membranes by enhancing membrane stiffness and lowering lipid fluidity. Moreover, the nanoshell serves as armor, protecting cells, limiting swelling and ultimately improving their viability against mechanical stress from osmotic imbalance and centrifugal forces. Our results demonstrate the potential of the nanoshell, not only as a cellular protection strategy, but also as a platform for manipulating and studying plasma membrane mechanics.

## Introduction

The cellular plasma membrane serves as a protective barrier by encapsulating cellular components.^1–3^ This biomembrane is decorated with membrane-bound proteins, making it essential for mediating cellular signaling and sensing.^1–4^ The plasma membrane is also linked to the interior cytoskeleton that mechanically supports the cell to maintain its size, shape and integrity.^5,6^ The cell membrane therefore allows for cellular communication while shielding the cell from outside assaults. However, the cell membrane is often unable to protect the cell from external stressors, for example, the high forces and subsequent large membrane deformations experienced during cell manipulation and delivery applications in tissue engineering and regenerative medicine.^7–9^

Cell encapsulation is recognized as one approach to tackle this problem with various nanomaterials being extensively investigated to wrap the whole cell for cellular protection and manipulation.^10–12^ However, the lack of material programmability limits the precision of the encapsulation, such as the control over the amount of membrane-deposited materials and their on-demand removal. Moreover, material overload as well as the cytotoxic nature of certain materials may hinder cell function and even lead to cell death.^13–16^ Recently, structural DNA nanotechnology including DNA origami has emerged as a powerful tool for studies interfacing cell biology with engineered nanostructures.^17–23^ A nanoshell approach that combines the DNA origami technique and cell membrane engineering, therefore has the potential to enable modular cell encapsulation that has the needed versatility, programmability, biocompatibility and biodegradability.^19,22^

In this work, we describe a nanoshell encapsulation strategy that targets the cell-surface glycocalyx, which utilizes two layers of DNA nanorods by sequentially recruiting and crosslinking them onto cell membranes under physiological conditions. We demonstrated the modularity and tunability of the nanoshell by varying the layering and composition of DNA nanorods. Although DNA nanostructures have been applied to encapsulated cells, previous biophysical and biomechanics studies have been performed primarily on artificial lipid bilayers coated with DNA nanostructures.^12,24–27^ The effects of such encapsulation on the live cell membrane biomechanics have not been fully explored. Further, the reconfiguration and polymerization of membrane-coated DNA origami on giant unilamellar vesicles (GUVs) may alter the coating pattern and induce membrane deformations, showing the dynamic interactions between the membrane and membrane-bound assemblies.^25,28–30^ However, live cell membranes are mechanically supported and protected by the cytoskeleton and the complex membrane-bound protein networks. Whether they will react similarly to DNA origami-coated GUVs remain unanswered. We therefore investigated the impact of the nanoshell on the biophysical properties of our cell-nanoshell systems by examining cell membrane biomechanics, membrane lipid fluidity and the distribution and morphology of DNA constructs after anchoring onto the membrane. Moreover, we demonstrated the protective effects of the nanoshell via enhanced viability after two environmental stressors: osmotic swelling and centrifugation. With this encapsulation strategy, we can build nanoshell potentially not only for cellular protection and ruggedization, but also as a tool for modulating membrane biomechanics and exploring the effects of these changes on cell behavior and function.

## Results and discussion

### Synthesis of DNA nanoshell on the cell membrane by crosslinking DNA nanorods

The nanoshell is designed to consist of two layers of crosslinking DNA nanorods that we referred to as rod A and rod B (both *∼* 7 nm in diameter and *∼* 400 nm in length). The rods were decorated with multiple functional ssDNA binding overhangs (Figure 1A and Supplementary Figure 1). Specifically, rod A had three anchoring ssDNA (a-ssDNA) for allowing the anchoring of rod A to membrane glycocalyx-anchored a-ssDNA complementary initiators (a’-ssDNA), 14 uniformly distributed staining ssDNA (s-ssDNA) for biotin attachment and subsequent streptavidin-fluorophore staining, and 14 uniformly distributed hybridization ssDNA (h-ssDNA) for the cross binding of rod A and rod B through ssDNA hybridization. Rod B nanostructures had 14 s-ssDNA and 14 hybridization ssDNA complementary (h’-ssDNA) that allowed them to bind to h-ssDNA-decorated rod A. We noticed that a higher number of h-ssDNA on rod A led to unstable rod formation (Supplementary Figure 2). An atomic force microscopy (AFM) image verified the formation of the rods, which were constructed as six-helical bundles (Figure 1A). Gel electrophoresis confirmed the monodispersity of the individual DNA origami rods. When combined, the formation of aggregate indicates the successful binding of the two rod species by hybridization (Figure 1B). Two other distinct bands suggest that rod mononers and dimers existed at the same time. As the nanorods will be used for cell culture applications, we further examined their stabilities in cell culture medium (Figure 1C). Individual rods had minimal degradation after 6 h and 30 h incubation at 37°C in both cell culture medium. The aggregate, formed by the combination of rod A and B, did not degrade noticeably for up to 30 h of incubation as well.

**Figure 1:**
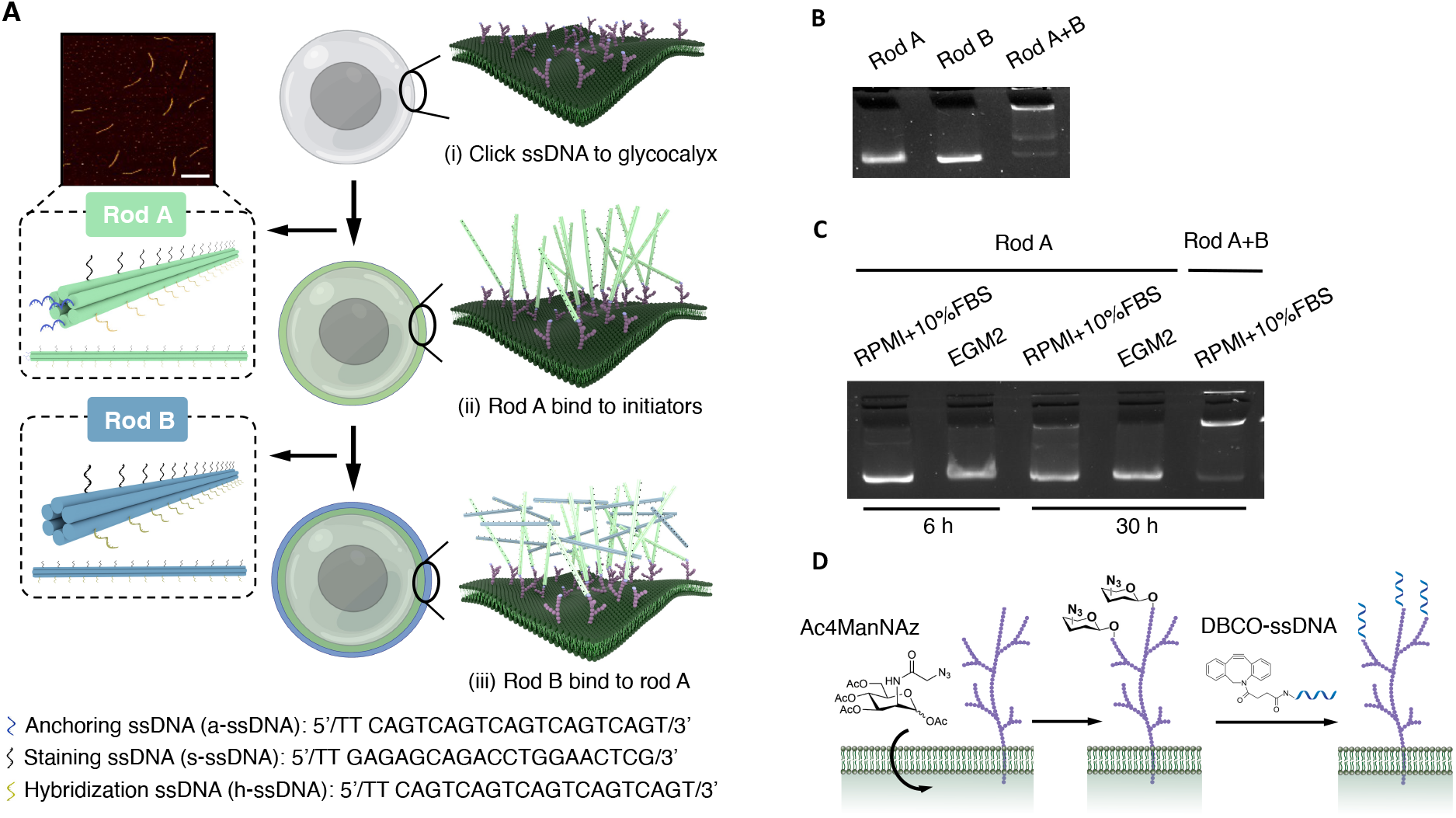
Schematics of the nanoshell synthesis process and validation of stability. (A) Synthesis of the DNA nanoshell on the cellular membrane through a three-step immobilization process including: (i) the immobilization of a’-ssDNA initiators on the glycocalyx; (ii) the binding of rod A (green) to a’-ssDNA via ssDNA hybridization and (iii) the binding and crosslinking of rod B (blue) to rod A via the hybridization of h-ssDNA on rod A and h’-ssDNA on rod B. AFM imaging of rods was shown. Both rod A and B were *∼*7 nm in diameter and *∼*400 nm in length. Three a-ssDNA (blue), 14 s-ssDNA (black) and 14 h-ssDNA (yellow) were uniformly distributed on rod A. 14 s-ssDNA (black) and 14 h’-ssDNA (yellow) were uniformly distributed on rod B. All ssDNA overhangs were 22 base pairs. Scale bar: 500 nm. (B) Agarose gel electrophoresis of individual DNA rods, and a mixture of rods after 30 min incubation of rod A and rod B at 37°C. (C) Agarose gel electrophoresis studies of the stability of individual DNA rods and the aggregate in two types of cell culture medium. Rod A and rods mixtures were incubated at 37°C for 6 h and 30 h in each cell culture medium. (D) The immobilization of DBCO-labeled a’-ssDNA initiators on azide-presenting cell-surface glycocalyx through copper free click chemistry.

To anchor rod A to the plasma membrane, using Jurkat cells as a suspended mammalian cell model, we utilized a method we have previously reported to first immobilized a’-ssDNA initiators onto the cell-surface glycocalyx (Figure 1D).^31^ In this method, azide ligands were covalently incorporated onto glycocalyx through metabolic glycan labeling using an azido monosaccharide, N-azidoacetylmannosamine-tetraacylated (Ac4ManNAz). a’-ssDNA were conjugated with dibenzocyclooctyne (DBCO) to form DBCO-a’-ssDNA through an NHS-Ester and amine reaction. Bioorthogonal glycocalyx labeling with copper-free click chemistry allowed the conjugation of azide ligands on glycocalyx and DBCO-a’-ssDNA, leading to the immobilization of a’-ssDNA on glycocalyx. For these studies, rod A was first introduced for glycocalyx binding, followed by Rod B for hybridization to immobilized rod A.

We observed the successful recruitment of rod A to the membrane and rod B to rod A using fluorescence microscopy (Figure 2A). Confocal microscopy cross-sectional images of cells coated with both rods and fluorescence intensity profiles extracted from those images further confirmed the binding of both rod A and rod B to the cell membrane and the formation of a nanoshell structure (Figure 2B). To confirm the efficacy of this approach on more than one mammalian cell type, we replicated the synthesis strategy on human umbilical vein endothelial cells (HUVECs), demonstrating the versatility and utility of this nanoshell encapsulation technique for both non-adherent and adherent cell types (Supplementary Figure 3). As the glycocalyx is on the surface of almost every mammalian cell, we expect this glycocalyx-targeting method to be applicable to a broad array of cell types.^32,33^

**Figure 2:**
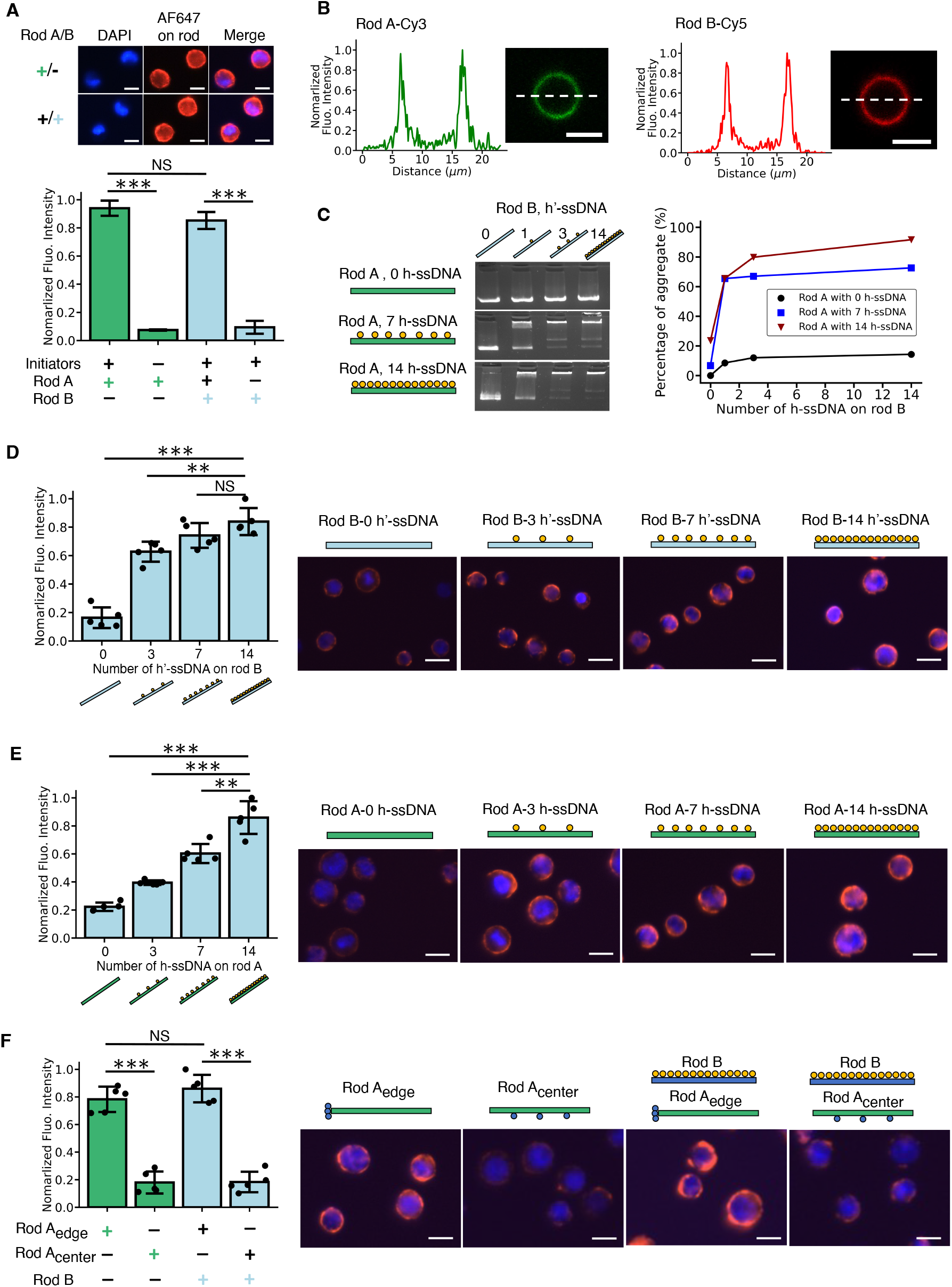
Modulation of the DNA nanoshell composition by engineering the position and number of overhangs on rods. (A) Fluorescence imaging and intensity quantification of four experimental conditions including: (i) rod A (biotin) coating with a’-ssDNA initiators on the glycocalyx; (ii) rod A (biotin) coating without a’-ssDNA initiators; (iii) rod B (biotin) coating with both the initiators and rod A, and (iv) rod B (biotin) coating with initiators but without rod A. Biotin-labeled rods were stained with streptavidin-AF647 (red). Cell nuclei were stained with DAPI (blue). The green and blue +/- symbols and accompanying bar graphs indicated whether rod A or rod B populations, respectively, were fluorescently labeled. (B) Normalized fluorescence intensity profiles of nanoshell-coated cells with Cy3-staining rod A (green) and Cy5-staining rod B (red). (C) Gel electrophoresis studies for the quantification of the percentage of aggegate resulting from each mixture of DNA rod A and B as a function of the number of hybridization overhangs on rods. The number of h-ssDNA on rod A was changed from 0, 7 to 14 and the number of h’-ssDNA on rod B was changed from 0, 1, 3 to 14. (D) Modulating the binding of rod B to rod A by changing the number of h’-ssDNA on rod B. The number of h’-ssDNA were changed from 0, 3, 7 to 14. Rod A with 14 h-ssDNA were used in this experiment. Rod B were stained with AF647 (red). (E) Modulating the binding of rod B to rod A by changing the number of h-ssDNA on rod A. The number of h-ssDNA were changed from 0, 3, 7 to 14. Rod B with 14 h’-ssDNA were used in this experiment. Rod B were stained with AF647 (red). (F) Modulating the binding of rod A onto the cell membrane and rod B to rod A by changing the position of a-ssDNA on rod A. The fluorescence intensity quantification of four conditions includes: (i) edge-decorated rod A (biotin); (ii) center-decorated rod A (biotin); (iii) edge-decorated rod A and rod B (biotin) and (iv) center-decorated rod A and rod B (biotin). Biotin-labeled rods were stained with streptavidin-AF647 (red). Schematics of rods and representative fluorescence microscope images were shown. Data were presented as means ± s.d. in (A), (D), (E) and (F). *\*\*P ≤* 0.01, *\*\*\*P ≤* 0.001. All scale bars: 10 *µ*m.

### Modulation of the DNA nanoshell composition

To demonstrate our ability to engineer the nanoshell, we systematically probed the roles of functional ssDNA binding overhangs extending from the DNA nanorods, investigating how the multivalency and positions of overhangs modulate the amount of rod binding in the nanoshell. First, we performed gel electrophoresis studies by mixing DNA rod A and B with varying number of h-ssDNA and h’-ssDNA in suspension and incubating for 0.5 h at 37°C to allow for binding. The number of evenly displayed h-ssDNA on rod A was modified from 0, 7 to 14, and the number of h’-ssDNA on rod B was from 0, 1, 3 to 14. Gel electrophoresis images and the quantification showed a monotonic decline of single rod band intensity and a monotonic increase in the aggregate band intensity with increasing number of h-ssDNA and h’-ssDNA (Figure 2C). To confirm this finding on Jurkat cells, we first labeled the cells with rod A bearing 14 h-ssDNA overhangs. Next, we introduced fluorescently labeled rod B with 0, 3, 7 and 14 h’-ssDNA. We found the binding of rod B increased monotonically again with increasing number of h’-ssDNA on rod B (Figure 2D). In addition, fluorescently labeled rod B with 14 h’-ssDNA were introduced to bind to rod A with 0, 3, 7 and 14 h-ssDNA. A similar trend of an increasing amount of rod B binding with increasing valency of h-ssDNA on rod A was observed (Figure 2E). Furthermore, the binding of rods was also increased by adding higher concentrations of rods (Supplementary Figure 4).

While we learned that the multivalency of hybridization ssDNA on rods A and rod B regulated the recruitment of rod B onto rod A, we also found that the position of anchoring ssDNA on rod A to be critical for the recruitment of both rods onto cell membranes. Rod A were modified to display a-ssDNA in two configurations: at the edge and at the center (Figure 2F). The binding of edge-decorated rod A to the glycocalyx and subsequent binding of rod B to rod A were significantly more than that of center-decorated rod A. This finding is consistent with previous studies, stating that the recruitment of DNA nanostructures presenting ssDNA overhangs at the sharp or “pointy” areas is more efficient.^23,34,35^ Our results demonstrate our ability to modulate the amount of nanorods incorporated into the nanoshell by changing the valency and positioning of functional ssDNA overhangs on both rods. As the maximum thickness of the nanoshell is defined by the length of rod A, an increase in rod binding indicates a higher density of rods. As a result of these findings, all following studies were performed with three edge-located a-ssDNA and 14 side-located h-ssDNA on rod A, and 14 side-located h’-ssDNA on rod B.

### Evaluation of nanoshell stability, morphology and rod distribution on cell membranes

Cells constantly internalize substances outside the membrane, inducing membrane remodeling and deformations at multiple scales.^36,37^ It is therefore important to evaluate the stability of the nanoshell on the membrane, which will be particularly instructive for future DNA nanostructure-based biomedical applications, for example, the longer-term presentation of functional nanodevices and biomolecules on nanoshells. We first evaluated surface retention time of the nanorods on the cell membrane as an indicator of stability. Nanoshell-coated cells and rod A-coated cells were incubated under three conditions: (i) 4°C for 0.5 h, (ii) 37°C for 0.5 h and (iii) 37°C for 3 h. The first incubation condition was regarded as a baseline as membrane movements and cell activities, especially cellular uptake, which lead to the destabilization of the nanoshells, were minimal. For consistent comparison, only rod A were stained with streptavidin-AF647 and they were stained after incubation. This staining method allowed us to only quantify the rods remaining on the external cell surface. From fluorescence-activated cell sorting (FACS) data, rod fluorescence intensity had a dramatic drop at 0.5 h incubation at 37°C as compared to 4°C, suggesting that single DNA rods had low stability after being anchored on cell membrane under physiological conditions, potentially due to cellular uptake or detachment from the membrane (Figure 3A). The fluorescence signal continued to decrease with further incubation at 37°C to 3 h, though at a slower rate. In contrast, on nanoshell-coated cells, we observed only a minimal decrease in fluorescence signal intensity with incubation at 37°C, even after 3 h (Figure 3A). This improved surface retention time of rods showed that the crosslinking nanoshell had a higher stability and remained on the cell membrane for a longer duration, as compared to single rod attachment without crosslinking. Moreover, although having a high stability under physiological conditions, the nanoshell can still be degraded through the simple administration of DNase I, making temporary encapsulation possible (Supplementary Figure 5). This feature is particularly important for applications requiring on-demand release or reconfiguration of the nanoshell.

**Figure 3:**
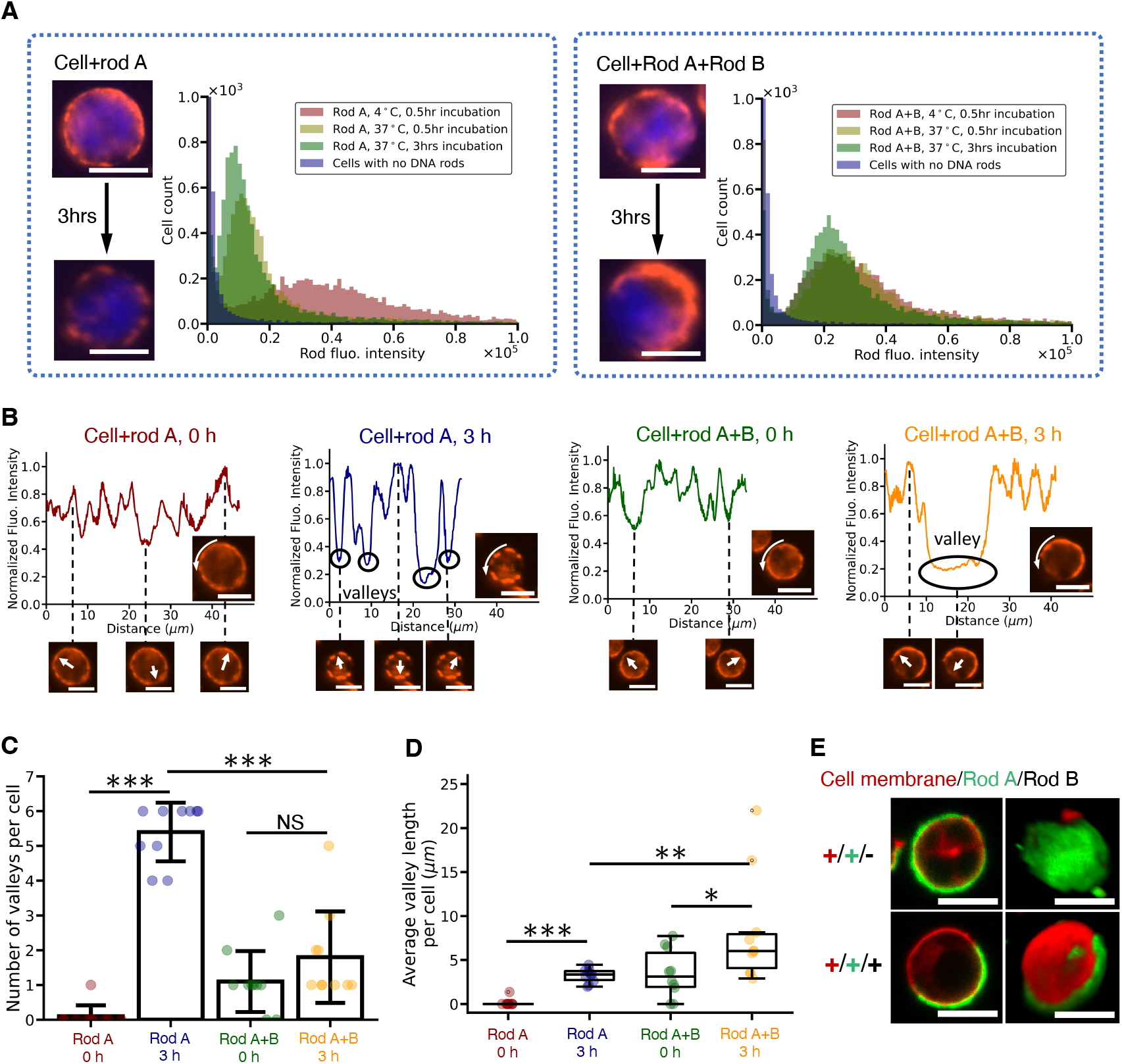
Cell surface retention time and morphology of the DNA nanoshell. (A) Cell surface retention time of DNA rods measured by fluorescence-activated cell sorting (FACS). Rod A-coated cells and nanoshell-coated cells were incubated at 4°C for 0.5 h and 37°C for both 0.5 h and 3 h. The distribution of cell populations versus AF647 fluorescence intensity were plotted. Representative wide-field fluorescence microscope images were taken. Pre-treatment native cells without fluorophore staining were used as control. DNA rod A were stained with streptavidin-AF647 (red). Cell nuclei were stained with DAPI (blue). (B) Analysis of DNA rod distribution on the borders of cell membranes. The fluorescence intensity along the cell borders was tracked and plotted for rod A-coated cells and nanoshell-coated cells after 0 h and 3 h incubation. DNA rod A were stained with AF647 (red). Wide-field fluorescence microscopy images showed the distribution of fluorescence signal on cells. Signal valleys were shown and marked in the plots. Signal valley quantification was reported in terms of (C) the number and (D) the length of fluorescence signal valleys in each group. *n* = 10. *\*P≤* 0.05, *\* *P≤* 0.01, *\* * * P≤* 0.001. (E) Confocal microscopy cross-section images and reconstructed 3D models of representative rod A-coated cells and nanoshell-coated cells. DNA rod A were stained with Alexa fluor 488 (green). Cell membranes were stained with DiD lipid dye (red). All scale bars: 10 *µ*m.

Next, we sought to understand whether the DNA rods and nanoshells interact with the membrane after rods were anchored onto the membrane. We observed substantial remodeling of nanoshell as the pattern of rod fluorescence signal evolved throughout incubation period (cell fluorescence images in Figure 3A). We tracked the distribution of fluorescence signals in cell images from these studies and measured the signal intensity around the contour of cell borders (Figure 3B). At 0 h time point where there were only minimal cell activities, relatively continuous and uniform intensity was observed in rod A-coated cells. Interestingly, however, the signal became discretized and non-uniform after 3 h incubation with signal intensity disappearing in discrete regions on cell borders, which resulted in signal valleys, suggesting long-term incubation destabilized the membrane-anchored DNA rods. Signal valleys also appeared in nanoshell-coated cells but were fewer in number and were substantially wider, potentially due to the crosslinking and polymerization of rods. We then quantified the number and the length of signal valleys (Figure 3C,D). Data showed that the discretization of fluorescence signals was dependent on two factors: incubation time and the addition of crosslinking rod B. Incubation-induced signal valleys were presumably due to cellular uptake whereas crosslinking-induced valleys suggested that the rods remodeled their positions on the cell membrane during crosslinking process and incubation. A similar phenomenon was also reported previously on GUVs where the distribution of DNA origami and the morphology of lipid bilayers were altered after the triggered polymerization of DNA origami.^28,29^ We further imaged rod A-coated cells and nanoshell-coated cells with confocal microscopy and reconstructed their three-dimensional models. The images revealed uniform covering on rod Acoated cells and partial, localized coverage on nanoshell-coated cells (Figure 3E). Our findings showcase the important role of the dynamic interactions between the DNA rods and the cell membrane in repositioning membrane-anchored substances. However, in this study, we did not observe membrane deformations due to DNA construct polymerization, which has been reported in the previuos literature.^25,28,29^

### Effects of nanoshell coating on cell membrane biomechanics

After demonstrating the ability of the nanoshell to remodel and stabilize itself, we sought to investigate whether this stabilization affects the biophysical properties of the cell membrane with a focus on membrane stiffness and lipid fluidity. First, membrane elastic modulus was evaluated by performing micropipette aspiration on pre-treatment native cells and nanoshell-coated cells (Figure 4A). Cells (*R ≈* 10 *µ*m) were aspirated into micropipettes (*R*_p_ =2.5 *µ*m) through aspiration pressure change Δ*P*. The elastic modulus *E* can be derived from the following equation, assuming the cell as a continuum-medium model with homogeneity,

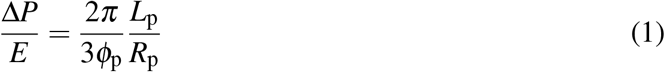

where *L*_p_ is the aspiration length and *φ*_p_ *≈* 2.1.^38,39^ The membrane elastic modulus of nanoshell-coated cells was 0.340 ± 0.062 *kPa* (mean ± s.d.), which was around 3-fold that of native cells, measured to be 0.122 ± 0.029 *kPa* (Figure 4B,C). Our results indicate that the nanoshell formed by crosslinking rods mechanically supported the membrane and enhanced the membrane mechanics, functioning analogously to the cytoskeleton underneath the membrane. Such an enhancement in membrane mechanics is consistent with crosslinking of our stiff 6-helix nanorod constructs. As we have demonstrated that we can engineer the density of the composites in the nanoshell by changing the valency and positioning of functional ssDNA overhangs on rods, we foresee that the degree of this mechanical enhancement may also be modulated accordingly.

**Figure 4:**
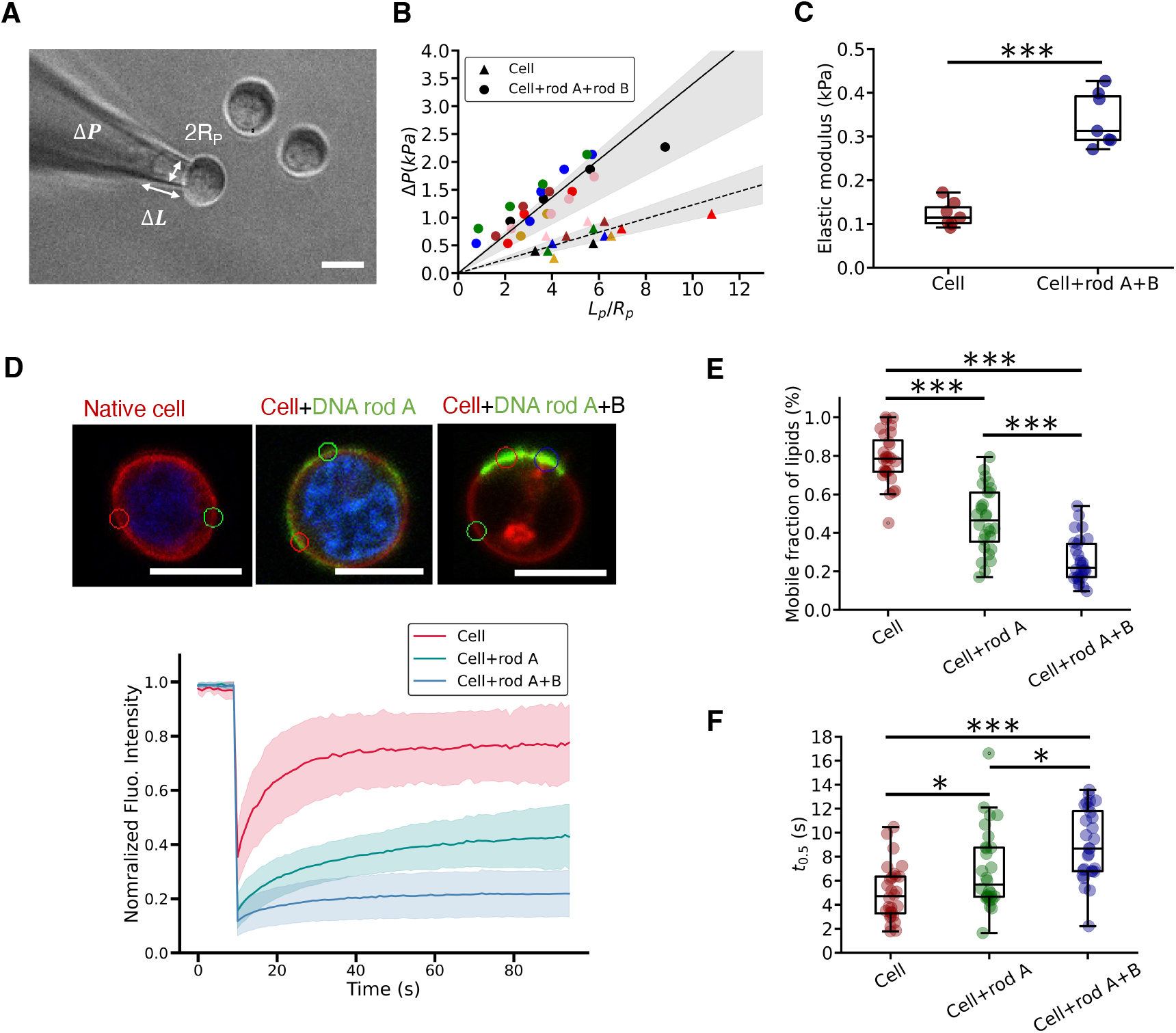
Direct modulation of biophysical properties of the cell by the DNA origami nanoshell. (A) A representative image of micropipette aspiration with a cell being aspirated into a micropipette with an inner radius *R*_p_ =2.5 *µ*m under an aspiration pressure Δ*P*. (B) Relationship between the Δ*P* and the normalized aspiration length 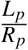. Triangular data: native cells; circular data: nanoshell-coated cells. The colors indicate different cell samples. (C) Evaluation of membrane elastic modulus of native cells and nanoshell-coated cells.^38^ *n* = 7 for (B) and (C). (D) FRAP experiments and the normalized membrane lipid fluorescence intensity over time after photobleaching a region of interest on the membranes of native cells, rod A-coated cells and nanoshell-coated cells. Data are presented as means (solid lines) ± s.d. (shadow areas) (*n* = 10). Cell membranes were stained with DiD lipid dye (red). DNA rod A were stained with streptavidin-AF488 (green). Evaluation of (E) of the mobile fraction of membrane lipid and (F) half-time recovery of membrane lipid of native cells, rod A-coated cells and nanoshell-coated cells. *n* = 28 for (D), (E) and (F). *\*P ≤* 0.05, *\*\*\*P ≤* 0.001. All scale bars: 10 *µ*m.

Given the global mechanical impact of the nanoshell on cell membrane mechanics, we sought to determine if the nanoshell could induce local changes to the fluidity of membrane lipid. A previous study found that by decorating DNA origami on GUVs with cholesterol anchors, the fluidity of artifical lipid was not affected.^26^ To address this question, we performed fluorescence recovery after photobleaching (FRAP) experiments to evaluate the mobile fraction of membrane lipid and the rate of recovery (Figure 4D). Interestingly, we observed that our glycocalyx-anchored nanoshell greatly reduced lipid mobility (Figure 4E). Specifically, in native cells, fluorescence signal recovered to *∼*80% of the pre-photobleaching level within 30 s. However, for rod A-coated cells, the recovery dropped to *∼* 40%. The number decreased further to *∼* 20% when the two-component nanoshell applied. The recovery here represents the mobile fraction of membrane lipid. We also noticed the rate of recovery was much slower after cells were coated with DNA rods (Figure 4F). The half time recovery of nanoshell-coated cell membrane lipid was only around half of that of native cell membrane lipid. These results demonstrate that a certain amount of membrane lipid experienced gelation when the membrane was coated with DNA rods and nanoshell, especially the latter case, resulting in a significant decrease in membrane lipid mobility. Intriguingly, the membrane-anchored DNA rods also lost their mobility whereas by comparison, DBCO-cy5 had nearly the same mobility as membrane lipid (Supplementary Figure 6). Taken together, these observations suggest that reductions in lipid mobility are not due to the metabolic glycan labeling nor the click conjugation of azide ligands and DBCO molecules, but rather due to the recruitment of DNA origami. Their large molecular weights and the potential spatial hinderance may be responsible for the low mobility of DNA origami. Our findings also show that the crosslinking of DNA rods further decreased the fluidity of membrane lipid, presumably due to enhanced rod-membrane interactions induced by rod crosslinking.

### Cellular protective effects of the nanoshell armor

The cell membrane is vital for maintaining cell size, shape and integrity, protecting the cell from outside assaults. The enhancement in membrane mechanics and the gelation of certain membrane lipid due to the nanoshell coating shows the potential of the nanoshell in providing protection to cells under harsh and mechanically challenging environments. To investigate its protective potential, we first examined the cell viability after cells were coated with nanoshell as a baseline. As expected, nanoshell-coated cells had high viability as the DNA nanostructures were biocompatible and the whole synthesis process was performed under physiological conditions (Supplementary Figure 7). We then applied osmotic imbalanced solutions to cells by changing the sodium chloride (NaCl) concentration from 0.9% to 0.6%, 0.3% and 0%. The resulting cell sizes and viability were measured (Figure 5A-C). As the osmotic pressure decreased, we found a rapid increase in the sizes of native cells and a decrease in their viability, whereas nanoshell-coated cells maintained the cell size and had a *∼* 20% higher cell viability in lower osmotic solutions, with statistical differences compared to pre-treatment native cells (Figure 5B,C). Notably, the nanoshell systems even maintained cell shape under 0% NaCl after 10 min incubation. In contrast, native cells and rod A-coated cells burst rapidly within seconds. Although the rod A coating was also able to limit cell expansion, it was not able rescue cell viability in low NaCl concentration solutions. We also noticed a slight decrease in the baseline cell size under 0.9% NaCl with 115±30 *µm*^2^ for native cells, 109±24 *µm*^2^ for rod A-coated cells and 102±20 *µm*^2^ for nanoshell-coated cells, with statistical difference between nanoshell-coated cells and rod A coated cells. These findings demonstrate the utility of coating DNA origami on the membrane to maintain cell size. Even a simple coating of rod A can limit cell expansion and improve survival, but the crosslinked nanoshell with both rods A and B is most effective in limiting expansion and importantly, acting as an armor and improving the survival of cells. Next, we applied centrifugal forces to cells and found that the nanoshell coating was able to rescue viability under 1500 g and 3000 g (Figure 5D).^12^ Interestingly, rod A coating was also able to improve cell viability against centrifugation, similar to its ability in limiting expansion under osmotic swelling. Our results show that DNA origami and nanoshell coating protect cells from mechanically challenging environments.

**Figure 5:**
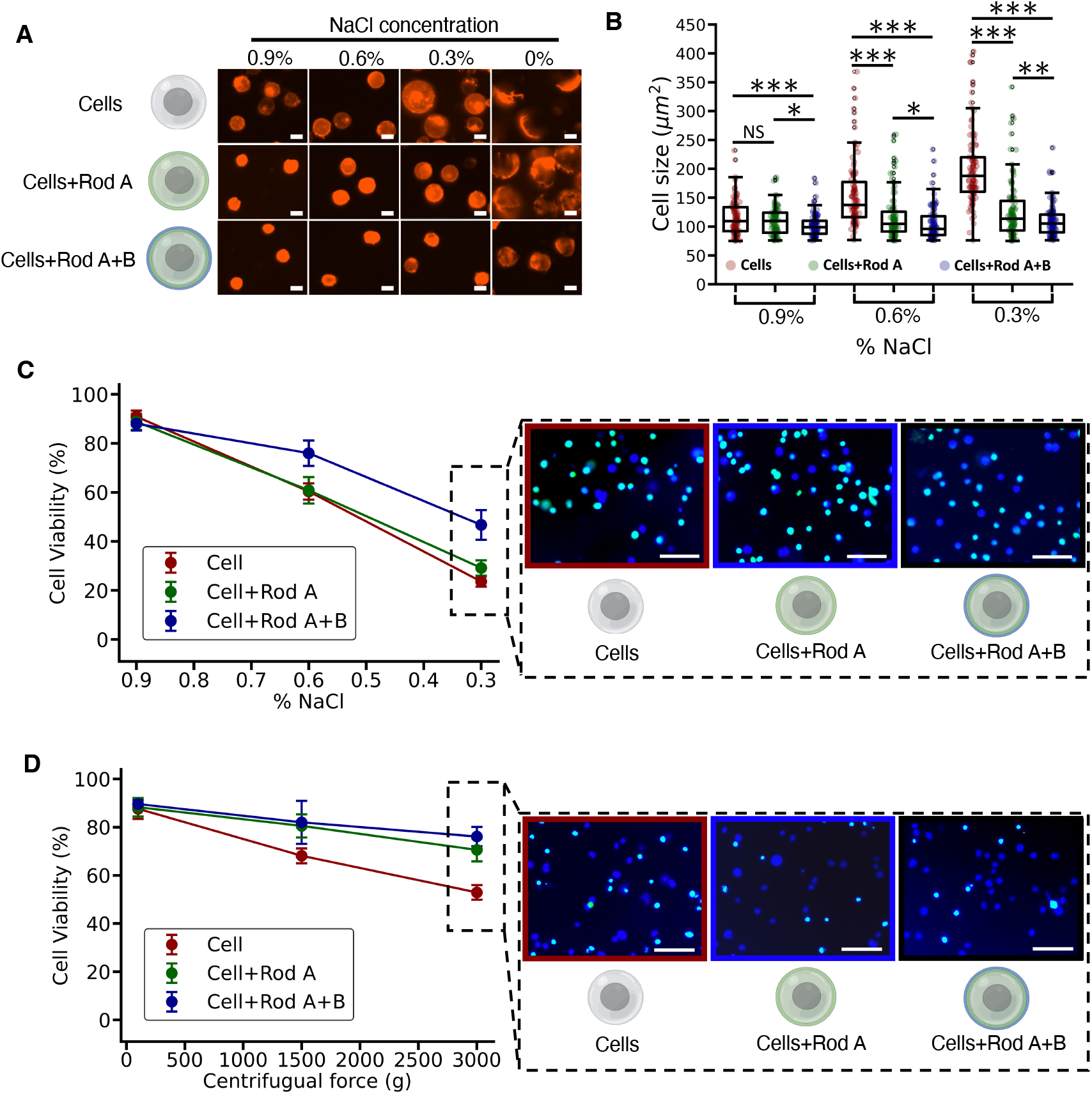
The protective effects of DNA nanoshell armor against challenging environments. (A) Fluorescence images of native cells, rod A-coated cells and nanoshell-coated cells after 10 min incubation in a range of concentrations (mass per unit volume) of NaCl from 0.9%, 0.6%, 0.3% to 0%. Cell membranes were stained with DiD lipid dye (red). Representative images were shown for each group. Scale bars: 10 *µ*m. (B) A bar plot for quantifying the cell size of native cells, rod A-coated cells and nanoshell-coated cells under osmotic imbalanced solutions. *n* = 100. (C) The relationship between cell viability and incubation in osmotic imbalanced solutions. Three conditions of NaCl concentration, 0.9%, 0.6% and 0.3%, were tested. Cells were LIVE/DEAD stained. Green: dead; blue: live. Representative cell images at 0.3% NaCl were shown. (D) The relationship between cell viability and centrifugation. A range of centrifugation rates were tested including 110 g, 1500 g and 3000 g. Cells were stained with the LIVE/DEAD staining after centrifugation. Representative cell images applied to 3000 g were shown. Scale bars for (C) and (D): 62.5 *µ*m. Data were presented as means ± s.d. as indicated by error bars (*n* = 3). Statistical tests were made to compare the viability of nanoshell-coated cells and native cells. *\*P ≤* 0.05, *\*\*P ≤* 0.01.

## Conclusions

In summary, we developed a modular strategy to encapsulate and ruggedize living cells using two layers of DNA origami nanorods. DNA rods were targeted and recruited to the cellular glycocalyx, followed by secondary rod crosslinking process, which successfully forms a biocompatible and biodegradable nanoshell under physiological conditions. We demonstrated that composition of the nanoshell was tunable by modifying the valency and position of overhangs on rods. We investigated multiple properties of the nanoshell, including the improved stability and the evolving migration of rod distribution triggered by incubation and rod crosslinking, thereby showing the impact of dynamic interactions within DNA rod assemblies as well as between rods and cell membranes. By probing the membrane biomechanics of nanoshell-encapsulated cells, we found that the nanoshell increased the membrane elastic modulus and inhibited the membrane lipid fluidity. Importantly, the nanoshell provided extra protection to cells, supporting enhanced viability despite harsh osmotic imbalance and centrifugal forces.

As DNA origami have been increasingly applied to cell biology, our study sheds light on the interactions between DNA nanostructures and cell plasma membrane, including the stability of the membrane-bound DNA nanostructures and the biophysical influences on cell membranes.^19–22^ Further, nanoshell-armored cells with viability enhancement may benefit biomedical and clinical applications such as regenerative medicine and cell printing.^10,12,14,40^

## Acknowledgement

This work was supported in part by the Air Force Office of Science Research grant number FA9550-18-1-0199 and a 2022 Dowd Fellowship. The authors thank Dr. Markus Deserno for membrane fluidity discussion, Molecular Biosensor and Imaging Center (MBIC) at Carnegie Mellon University (CMU) and Dr. Lydia A. Perkins for assistance on confocal microscope imaging, Mellon Institute for assistance on flow cytometry, Vismaya Walawalkar and Defense University Research Instrumentation Program (DURIP) for accessing AFM, and the Department of Chemical Engineering at CMU for access to the micropipette aspiration equipment.

