## Supplementary for "Synthetic cell armor made of DNA origami"

List of Supplementary Information:

Supplementary Figure 1. Cadnano design of DNA nanorod.

Supplementary Figure 2. Gel electrophoresis studies of rod A formation with varying number of hybridization ssDNA (h-ssDNA).

Supplementary Figure 3. DNA nanorod and nanoshell coating of adherent HUVECs.

Supplementary Figure 4. Modulation of nanoshell composition by the concentrations of DNA rods.

Supplementary Figure 5. DNA nanorods and DNA nanoshell degradation by DNase I.

Supplementary Figure 6. Fluorescence recovery after photobleaching (FRAP) experiments of native cells, rod A-coated cells and nanoshell-coated cells.

Supplementary Figure 7. Cell viability of native cells in control buffers and cells after DNA rod and nanoshell coating.

Table S1. List of functional DNA oligos.

Table S2. List of DNA oligos for DNA nanorods.

Materials and Methods

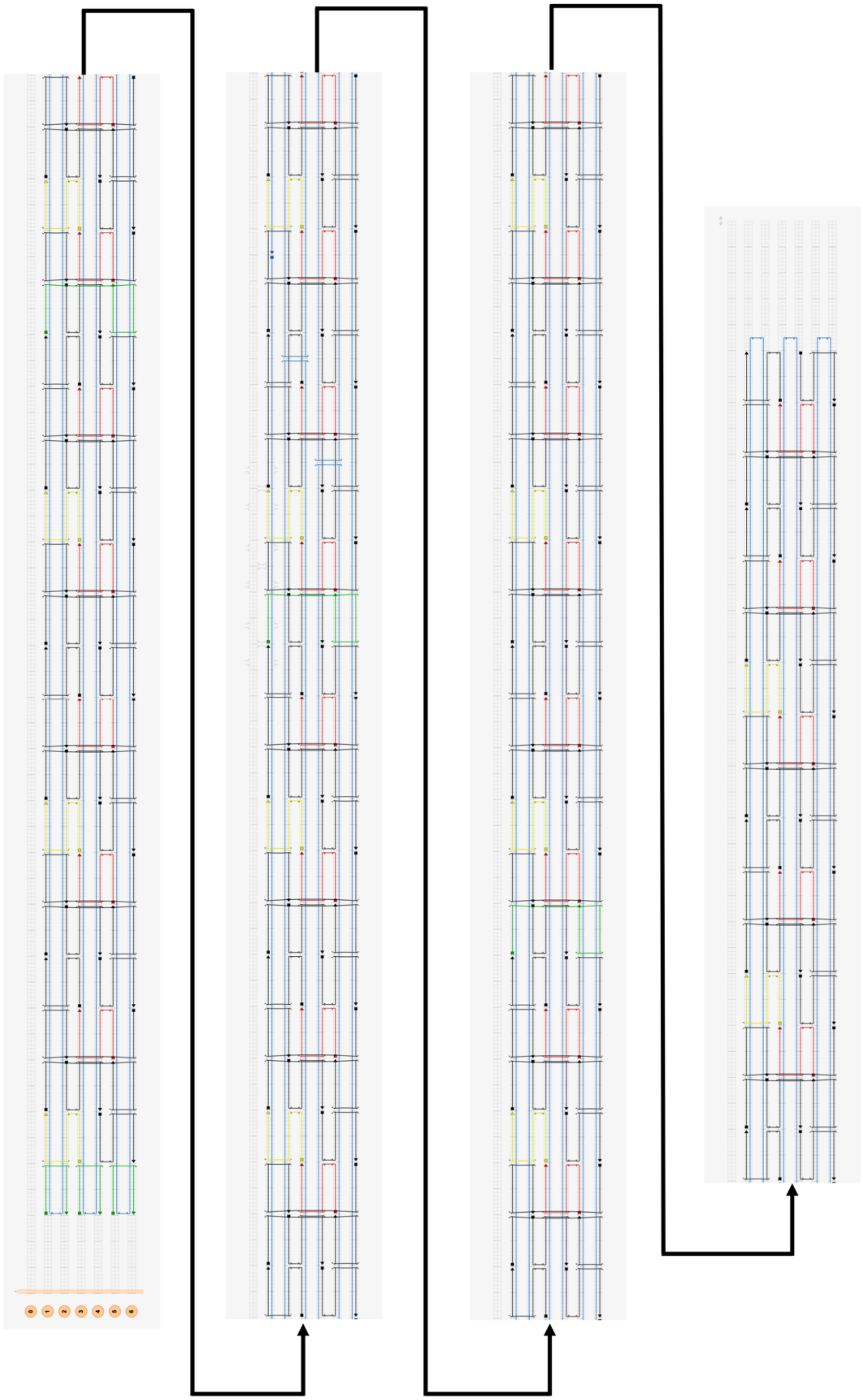

Supplementary Figure 1. Cadnano design of DNA nanorod. Green staples are modified with anchoring-ssDNA for rod A (rod B doesn't have green staples), 6 in total (3 at the edge are edge-decorated rod A. 3 in the center for center-decorated rod A). Yellow staples are modified with staining-ssDNA, 14 in total. Red staples are modified with hybridization-ssDNA for rod A and hybridization-ssDNA complementary for rod B, 28 potential sites in total. The naming of these sites is h-1 to h-28, from left to right.

| Lane | 1 | 2 | 3 | 4 |
|---|---|---|---|---|
| a-ssDNA | 3 | 3 | 3 | 3 |
| s-ssDNA | 14 | 14 | 14 | 14 |
| h-ssDNA | 7 | 14 | 21 | 28 |

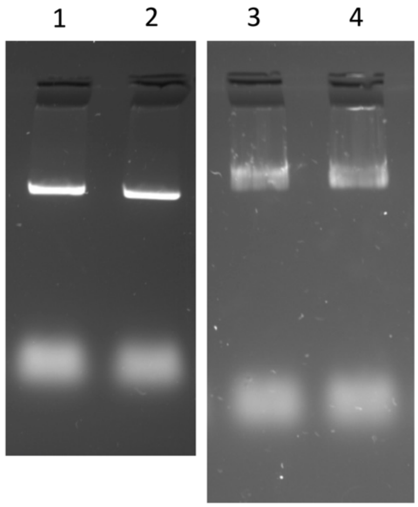


Supplementary Figure 2. Gel electrophoresis studies of rod A formation with varying number of hybridization ssDNA (h-ssDNA). Rod A were modified with 3 anchoring ssDNA, 14 staining ssDNA, and 7 or 14 or 21 or 28 h-ssDNA. Rod A had monodispersed individual bands when rod A was modified with 7 or 14 h-ssDNA, showing the formation of DNA origami rods. When modified with 21 or 28 h-ssDNA, rod A formation was not clear.

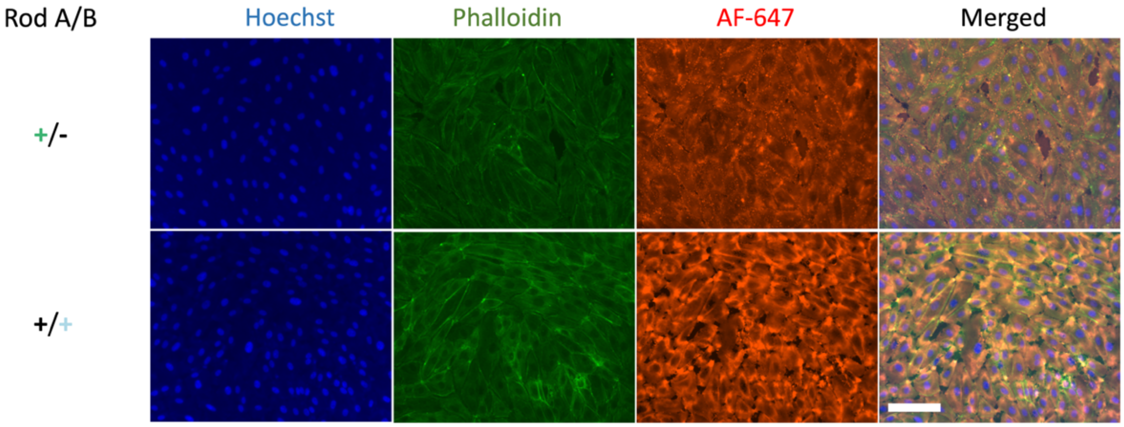


Supplementary Figure 3. DNA nanorod and nanoshell coating of adherent HUVECs. Cell nucleus were stained with Hoechst (blue). Cell membrane F-actin were stained with phalloidin (green). DNA nanorods were stained with AF-647. The experiment in the first row is cell + DNA rod A where rod A were stained. The experiment in the second row is cell + DNA rod A + rod B where rod B were stained. Scale bars: 125 µm.

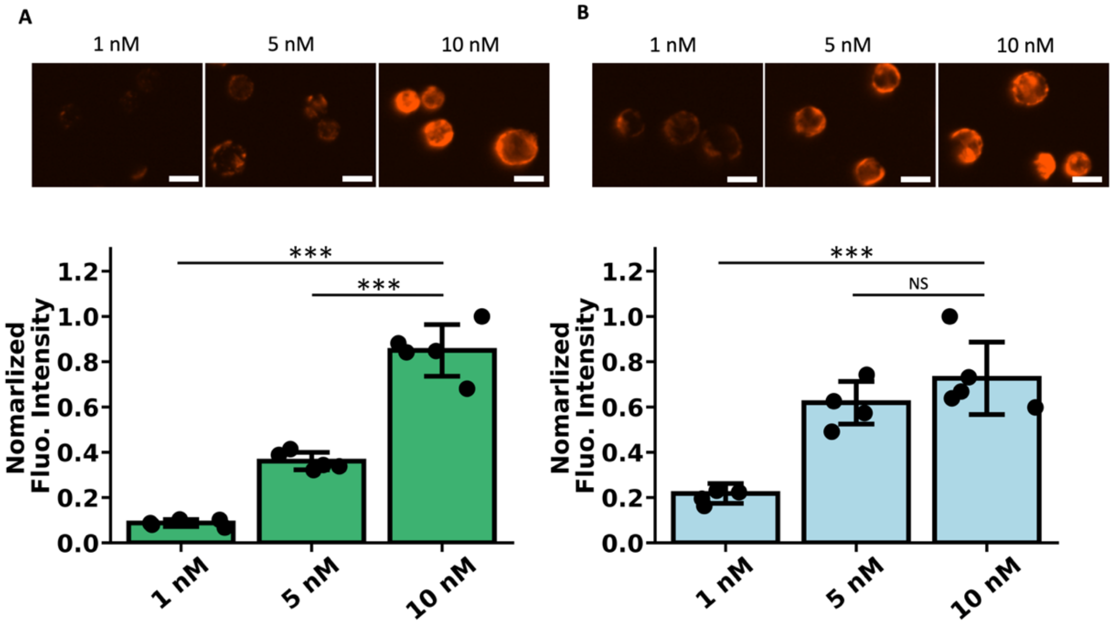


Supplementary Figure 4. Modulation of nanoshell composition by the concentrations of DNA rods. (A) The concentration of rod A was adjusted from 1 nM, 5 nM to 10 nM. The binding of rod A was measured by rod A fluorescence intensity (labeled with AF-647). (B) 5 nM of rod A were firstly administrated to cells. Then rod B were introduced where the concentration of rod B was adjusted from 1 nM, 5 nM to 10 nM. The binding of rod B was measured by rod B fluorescence intensity (labeled with AF-647). In both experiments, the volume of rods was 100 µl. Representative fluorescence microscopy images were taken. ***$P \leq 0.001$ (n=5). All scale bars: 10 µm.

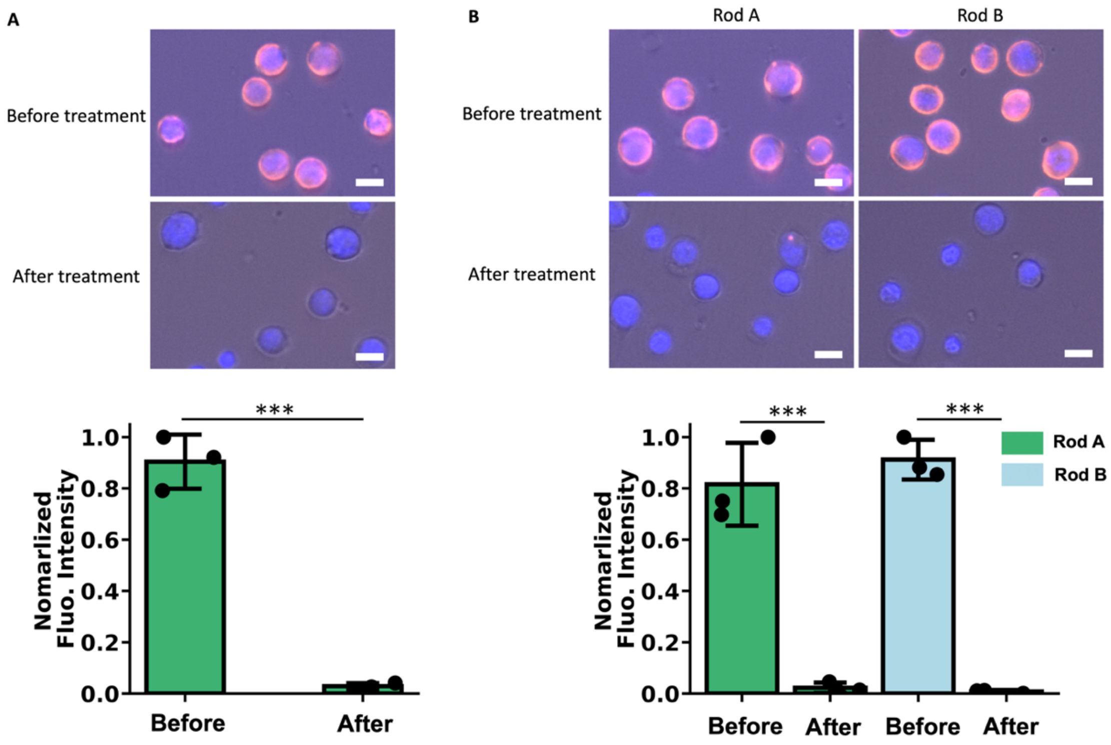


Supplementary Figure 5. DNA nanorods and DNA nanoshell degradation by DNase I. (A) Normalized fluorescence signal intensity of rod A on rod A-coated cells before and after DNase I treatment. (B) Normalized fluorescence signal intensity of rod A and rod B on nanoshell-coated cells before and after DNase I treatment. In both experiments, cell nucleus were stained with DAPI (blue). DNA rod A were stained with AF-647 (red). Representative bright-field and fluorescence images were shown. ***$P \leq 0.001$. All scale bars: 10 µm.

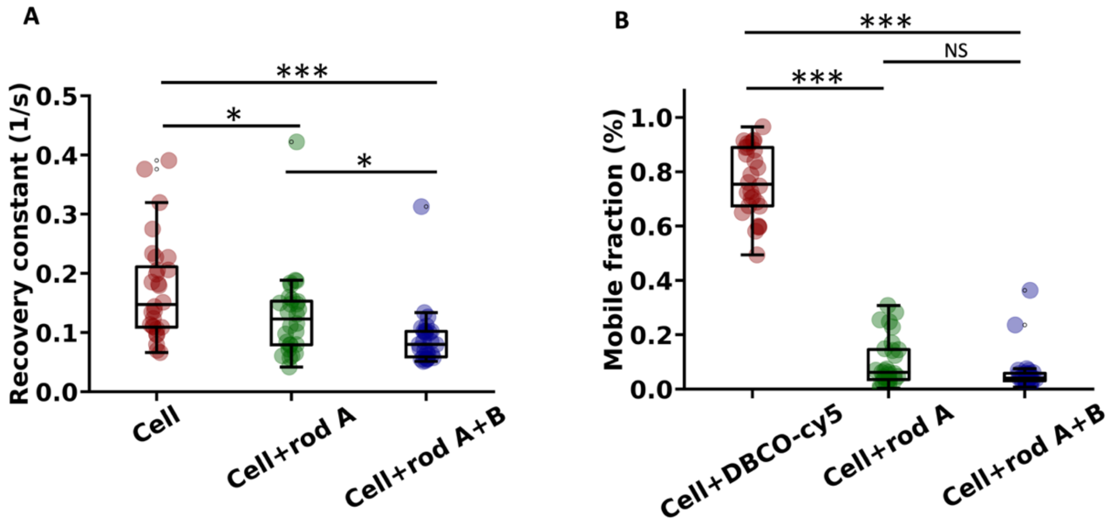


Supplementary Figure 6. Fluorescence recovery after photobleaching (FRAP) experiments of native cells, rod A-coated cells and nanoshell-coated cells. (A) recovery constant of cell membrane lipids and (B) the mobility of DBCO-cy5 in native cell+DBCO-cy5, the mobility of rod A in rod A-coated cells and the mobility of rod A in nanoshell-coated cells. n=28. *P ≤ 0.05, ***P ≤ 0.001.

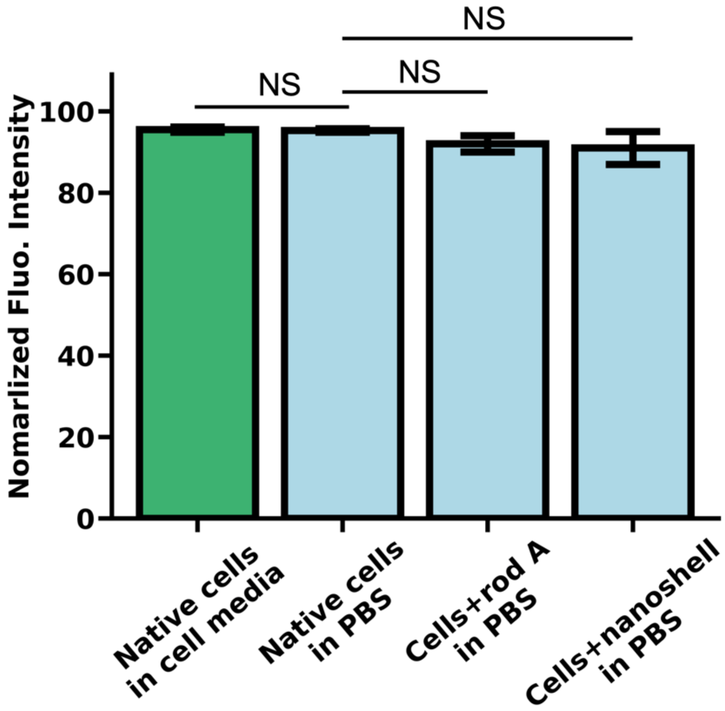


Supplementary Figure 7. Cell viability of native cells in control buffers and cells after DNA rod and nanoshell coating. Four conditions include native cells in cell media (RPMI+10%FBS), PBS and cells decorated with rod A and the nanoshell, both in PBS. Cell viability was assessed by ReadyProbes Cell viability imaging kit. Data are presented as mean ± standard deviation as indicated by error bars. n=3.

### Materials and Methods

#### Materials, reagents and equipment

DNA oligos (Table. S1) were purchased from Integrated DNA Technologies (Coralville, IA). DBCO-cy5 and DBCO-Sulfo-NHS-Ester, Sodium chloride and paraformaldehyde and PEG 8000 were purchased from Sigma-Aldrich. Streptavidin-Alexa Fluor 488 and 647 conjugates were purchased from Invitrogen. Roswell park memorial institute (RPMI) 1640 medium, dulbecco's phosphate-buffered saline (DPBS) was purchased from Corning. Bovine serum albumin (BSA) was purchased from Fisher BioReagents. AFM tips were purchased form NanoAndMore. Micropipettes (5 μm in diameter) were purchased from World Precision Instruments. Membrane lipid dye DiD and DNase I were purchased from ThermoFisher. ReadyProbes Cell viability imaging kit (Blue/Green) and SYBR Safe DNA gel stain were purchased from Invitrogen. Endothelial cell growth medium-2 bulletKit was purchased from Lonza.

#### Synthesis and purification of biotinylated DNA nanorods

DNA nanorods were folded from M13mp18 scaffold (Bayou Biolabs) together with 173 staple strands (rod A) and 170 staples (rod B). For annealing, 20 nM of the scaffold and 100 nM of the staples were mixed in TAE buffer with 12.5 mM $MgCl_2$. The biotin-labeled strands were added pre-annealing with the same concentration as other staples. The mixture was heated to 80°C for 5 min and gradually cooled down to 4℃ within 45 h. specifically, 80℃-65℃, 0.1℃ per 24 seconds; 65℃-24℃, 0.1°C per 378 seconds; 24°C-4°C, 0.1°C per 18 seconds. After annealing, the solutions were collected and precipitated by centrifugation at 10500 g for 25 min in a buffer containing 7.5% PEG 8000, 10 mM $MgCl_2$, 255 mM NaCl, 22.5 mM Tris, 10 mM acetic acid, and 1mM EDTA. The supernatant was removed and the pellet was resuspended to 5 nM using DPBS buffer with 12.5 mM $MgCl_2$. DNA nanorods were then administrated to cells within 30 min after purification.

#### Agarose gel electrophoresis and stability test

10 μl of 10 nM purified DNA nanorod samples were analyzed by electrophoresis in 2% agarose gel in TBE buffer with 12.5 mM $MgCl_2$ at 90 V and room temperature (RT) for 1 h. To study the structural stability of DNA nanorods in cell media, DNA rod A were

incubated in EGM-2, RPMI+10% FBS at 37°C for 6 h and 30 h before running the gel. To test the stability of rod A and B aggregate stability, rod A and B were resuspended in RPMI+10% FBS at 20 nM and were mixed and incubated at 37°C for 30 min and 30 h before running the gel. The gels were stained with 1×SYBR Safe DNA gel stain and imaged with Biorad ChemiDoc Imaging System.

Atomic force microscopy (AFM)

10 µl of 0.5 nM of purified DNA nanorod samples in DPBS with 12.5 mM $MgCl_2$ was deposited onto a freshly cleaved mica surface, incubated at RT in a humid chamber for 5 min. The sample was washed three times with 20 µl DI water and thoroughly blow-dried with nitrogen after each washing. AFM scans were performed using MFP-3D-BIO AFM (Asylum Research) with a 5 nM AFM tip in tapping mode.

Synthesis of 5'DBCO-a'-ssDNA

The 5'$NH_2$-a'-ssDNA oligos were incubated overnight in DPBS with DBCO-sulfo-n-hydroxysuccinimidyl ester (DBCO-Sulfo-NHS-Ester) at a 1:10 molar ratio under agitation at RT for 30 min. After incubation, the reaction mixture was dialyzed five times against DPBS using Amicon Ultra Centrifugal filters (molecular weight cut-off, 3 kDa) to remove unconjugated DBCO-Sulfo-NHS-Ester. The above-described conjugation and purification procedures were repeated twice in total to increase the conjugation efficiency. The final concentration of 5'DBCO-a'-ssDNA was adjusted to 500 µM with DPBS.

Cell culture

Jurkat, clone E6-1, were purchased from ATCC. Cells were cultured in RPMI+10% FBS at 37°C with 5% $CO_2$ and were maintained at a cell density between 100,000 cells/ml and 3,000,000 cells/ml. Human umbilical vein endothelial cells (HUVECs) were purchased from the Lonza. Cells were cultured in EGM-2 at 37°C with 5% $CO_2$. HUVECs of passage 3-5 were plated into 96-well culture plates at a density of 40,000 cells per well, one day prior to the experiment. Both cells have been tested negative for mycoplasma contamination.

Synthesis of DNA nanoshell on membrane through click chemistry and ssDNA hybridization

In the first step, we immobilized a'-ssDNA on glycocalyx. Azide ligands were firstly covalently incorporated onto glycocalyx 24 h before experiment, through metabolic glycan labeling using an azido monosaccharide, N-azidoacetylmannosamine-tetraacylated (Ac4-ManNAz). Specifically, Ac4ManNAz was diluted in culture medium to a final concentration of 50 µM and administered to cells 24 h before experiment. DBCO-a'-ssDNA was administrated to Jurkat cells at a final concentration of 50 µM and 100 µl in volume and incubated for 1 h. Cells were then washed to remove excess DBCO-a'-ssDNA using centrifugation at 200 g for 5 min. The washing step was repeated three times using DPBS+0.1% BSA as washing buffer. Next, 100 µl of 5 nM DNA rod A in DPBS with 12.5 mM $MgCl_2$ were administrated to cells and incubated for 30 min. Washing was performed three times after incubation. Then 100 µl of 5 nM DNA rod B in DPBS with 12.5 mM $MgCl_2$ were administrated to cells and incubated for 30 min. Cells then were washed three times and resuspended in DPBS or RPMI+10% FBS buffers for later experiments.

Examination of nanoshell cell surface retention time with flow cytometry

In this experiment, only DNA rod A were biotinylated and fluorescently labeled. Cells decorated with only rod A and with both rods A and B were incubated at three conditions: 4°C for 30 min, (ii) 37°C for 30 min and (iii) 37°C for 3 h, with RPMI1640+10% FBS as buffer. After incubation, cells were collected and fixed using 1 ml of 4% of paraformaldehyde with 10 min at RT. DPBS was used to wash for three times. DNA rod A were then stained with streptavidin-AF647 and cells were washed three times with DPBS. Finally, cells were analyzed using flow cytometry and fluorescence intensity of AF647 was examined by FACS.

Degradation of DNA nanorods and nanoshells on cell membranes

DNA rod A-coated cells and nanoshell-coated cells (1,000,000/ml) were incubated at 37°C for 30 min with a concentration of DNase I at 20 U and with DPBS as buffer. After incubation, cells were washed three times and imaged.

Quantification of fluorescence signal distribution on cell border

The fluorescence signals on cell borders were acquired using National Institutes of Health Image J by manually circling cell borders and recording the intensity.[1] The data were then analyzed by a custom Python 3.8 code using SciPy where the signal valley was defined as a fluorescence signal that was lower than 0.4, and the number and length of signal valleys were quantified.[2]

Examination of membrane lipid fluidity with FRAP

Live native cells, rod A-coated cells and nanoshell-coated cells were observed under confocal laser scanning microscope (LSM 880 with Airyscan, ZEISS). Cell membranes were stained with DiD lipid dye (red) and DNA rod A were stained with AF488 (green). Region of interest (ROI) was selected and set to be photobleached. The pinhole size for photobleaching was 2 μm. After photobleaching, the relation of fluorescence recovery with time was recorded and analyzed to calculate the mobile fraction, $t_{0.5}$ and recovery constant by ZEN (ZEISS).

Examination of cell membrane elastic modulus with micropipette aspiration

Live native cells and nanoshell-coated cells were deformed by glass micropipettes. Specifically, the micropipette was connected to tubing that ran to a water-filled reservoir with controllable height. The tip of the micropipette was then placed near a cell and a suction pressure was applied within the micropipette. The pressure difference deformed the cell and a membrane protrusion was formed. The length of the protrusion, the aspirated length, was recorded. Pressure difference and aspirated length were fitted linearly by forcing the fitted line to pass through the origin. Elastic modulus was calculated using the model and equation by Theret *et al.*, assuming the cell as a continuum-medium model with homogeneity.[3]

Protective effect of the nanoshell against osmotic swelling and centrifugal forces

In the osmotic swelling assay, native cells, rod A-coated cells and nanoshell-coated cells were diluted to a concentration of 1,000,000/ml. Cell membranes were stained by DID lipid dye (red). Cells were then centrifuged and resuspended in 1 ml of NaCl solutions (0%,

0.3%, 0.6%, 0.9%). After 10 min incubation at 37°C, cells were collected and imaged by a fluorescence microscope (EVOS). The sizes of cells were analyzed using Image J software by applying the same threshold to red areas in all images. To test cell viability, when incubating cells, NaCl buffers were mixed with cell viability imaging kit solutions. After incubation, cells were imaged on EVOS at 40x in both Cy5 and GFP channels. In the centrifugal assay, cells (1,000,000/ml) were centrifuged (centrifuge 5804R, eppendorf) five times at 4°C with a speed of 100 g, 1500 g and 3000 g, respectively. After each centrifugation, cells were resuspended by 1 ml of DPBS. Cells were then incubated in cell viability imaging kit solutions at 37°C for 10 min, followed by fluorescence microscopy imaging. Dead cells were stained green. All cells were stained blue. Cell viability was calculated as the ratio of live cell/all cells.

### Statistics and Reproducibility

Quantitative data were displayed as means ± s.d. Statistical significances were determined using one-way analysis of variance (ANOVA) with post-hoc Tukey's Test. All cell experiments were repeated independently at least three times.

Table S1. List of functional DNA oligos.

| Name of oligos | Sequence |
|---|---|
| Anchoring-ssDNA (a-ssDNA) modifier | 5'/TT CAGTCAGTCAGTCAGTCAGT/3' |
| staining-ssDNA (s-ssDNA) modifier | 5'/TT GAGAGCAGACCTGGAACTCG/3' |
| hybridization-ssDNA modifier (h-ssDNA, for rod A) | 5'/TT GTGACATACCTTCGGAGCAT/3' |
| hybridization-ssDNA complementary modifier (h'-ssDNA, for rod B) | 5'/TT ATGCTCCGAAGGTATGTCAC/3' |
| 5'$NH_2$-a'-ssDNA | /5AmMC6/TT ACTGACTGACTGACTGACTG |
| Biotin-s'-ssDNA | /5BiosG/TT CGAGTTCCAGGTCTGCTCTC |
| Cy5-s'-ssDNA modifier | /5Cy5/TT CGAGTTCCAGGTCTGCTCTC |
| Cy3-s'-ssDNA modifier | /5Cy3/TT CGAGTTCCAGGTCTGCTCTC |

### Table S2. List of DNA oligos for DNA nanorods.

| color in cadnano file | sequence | +a-ssDNA | +s-ssDNA | +1 h/h'-ssDNA | +3 h/h'-ssDNA | +7 h/h'-ssDNA | +14 h/h'-ssDNA |
|---|---|---|---|---|---|---|---|
| black | TCAGGCTGCGCAACCTAGGGCGCTGGCAATCGTCTGAAATGG | | | | | | |
| black | CATAACGCCAAAAGTTGCTAAACAACTTCCAATAGGAACCCA | | | | | | |
| black | CCGCTTCTGGTGCCCCACACCCGCCGCGACAGGAAAAACGCT | | | | | | |
| black | TATCGGCCTCAGGAATGGTTGCTTTGACTTGCTGGTAATATC | | | | | | |
| black | CATCGTAACCGTGCGAATCAGAGCGGGAATAACATCACTTGC | | | | | | |
| black | GATTGACCGTAATGTTAGACAGGAACGGTCACGCAAATTAAC | | | | | | |
| black | TCAGTTGAGATTTAAAGGAACAACTAAACCACCCTCAGAGCC | | | | | | |
| black | AACGAACTAACGGATGAAAATCTCCAAAGGTTTAGTACCGCC | | | | | | |
| black | TATACCAGTCAGGAGTATCGGTTTATCAATATAAGTATAGCC | | | | | | |
| black | ATCATTGTGAATTAAGCTTGATACCGATTTTTGCTCAGTACC | | | | | | |
| black | CGAGTAGTAAATTGGCCCACGCATAACCAGAGGCTGAGACTC | | | | | | |
| black | TCATTCAGTGAATAGAGTTAAAGGCCGCTGCCTATTTCGGAA | | | | | | |
| black | AGAACCGGATATTCAAAGACAGCATCGGGTGCCTTGAGTAAC | | | | | | |
| black | GGCGCATAGGCTGGTTGAGGACTAAAGAGATGATACAGGAGT | | | | | | |
| black | TGACCAACTTTGAAGGGTAAAATACGTATCTCTGAATTTACC | | | | | | |
| black | GCCGGAACGAGGCGCGAAAGAGGCAAAACAAACAAATAAATC | | | | | | |
| black | GATAAATTGTGTCGCCCAGCGATTATACGAAGTATGTTGAGG | | | | | | |
| black | TTGCGTATTGGGCGCTTTTCACCAGTGAAATAGATTAGAGCC | | | | | | |
| black | TATCATAACCCTCGCGTCTTTCCAGACGGTACAAACTACAAC | | | | | | |
| black | CAGCTGCATTAATGGCCTGGCCCTGAGATGAGGAAGGTTATC | | | | | | |
| black | TTGCGCTCACTGCCTGCCCCAGCAGGCGATCAATATCTGGTC | | | | | | |
| black | GCCTGGGGTGCCTATCGGCAAAATCCCTTCTAAAGCATCACC | | | | | | |
| black | ACAATTCCACACAAGGTTGAGTGTTGTTCCTGCAACAGTGCC | | | | | | |
| black | TCATGGTCATAGCTAGAACGTGGACTCCGCAGAAGATAAAAC | | | | | | |
| black | TCGACTCTAGAGGAAGGGCGATGGCCCAAGCCCTAAAACATC | | | | | | |
| black | GTTGTAAAACGACGTTTTGGGGTCGAGGAATATTTTTGAATG | | | | | | |
| black | GGCGATTAAGTTGGAAGGGAGCCCCCGAGAACCCTTCTGACC | | | | | | |
| black | CTTCGCTATTACGCACGTGGCGAGAAAGACACGACCAGTAAT | | | | | | |
| black | GAGAAGTGTTTTTAGTCGGATTCTCCGTAAATGTGAGCGAGT | | | | | | |
| black | ATTATTTACATTGGAATTAATTACATTTCGTTATACAAATTC | | | | | | |
| black | CATGGAAATACCTAATGGAAACAGTACACGGAATCATAATTA | | | | | | |
| black | CAGAACAATATTACTGCTTCTGTAAATCACCGACCGTGTGAT | | | | | | |
| black | CTGAGTAGAAGAACTCCTTGAAAACATATAGTTAATTTCATC | | | | | | |
| black | CGTTGTAGCAATACGAGTCAATAGTGAATCGCAAGACAAAGA | | | | | | |
| black | GAGGCCACCGAGTAACCTTTTTAACCTCGTTGGGTTATATAA | | | | | | |
| black | ACCACCCTCATTTTCAAAGACAAAAGGGAACAAAGTTACCAG | | | | | | |
| black | ACCCTCAGAACCGCGTAAATATTGACGGATCTTACCGAAGCC | | | | | | |
| black | CGGAATAGGTGTATCGTCACCGACTTGAAGCCCAATAATAAG | | | | | | |
| black | AGGCGGATAAGTGCCACCAGTAGCACCATGAGCGCTAATATC | | | | | | |
| black | CTCAAGAGAAGGATCAATGAAACCATCGGGGAGAATTAACTG | | | | | | |
| black | CCTATTATTCTGAAAATCAAGTTTGCCTTTTACAGAGAGAAT | | | | | | |
| black | AGTGCCCGTATAAACGGCATTTTCGGTCGAAACGATTTTTTG | | | | | | |
| black | GTACTGGTAATAAGTTTCATAATCAAAATTACAAAATAAACA | | | | | | |
| black | GTTCCAGTAAGCGTCGCCTCCCTCAGAGTACCAACGCTAACG | | | | | | |
| black | GCCTGTAGCATTCCCAACATATAAAAGAGCAGTATGTTAGCA | | | | | | |
| black | CTCATTAAAGCCAGAGCCACCACCCTCATAGTTGCTATTTTG | | | | | | |
| black | CAGGTCAGACGATTCGCCGCCAGCATTGACCTCCCGACTTGC | | | | | | |
| black | GTCAATAGATAATACAACTCGTATTAAAAAGGCTTATCCGGT | | | | | | |
| black | TAAAATATCTTTAGAAGTTTGAGTAACAAGGAATCATTACCG | | | | | | |
| black | AGTTGGCAAATCAACAGAAGGAGCGGAACGCACTCATCGAGA | | | | | | |
| black | TTGCTGAACCTCAAATGGCAATTCATCAGTCTTTCCTTATCA | | | | | | |
| black | ACGCTGAGAGCCAGTCTGAATAATGGAAATCCTAATTTACGA | | | | | | |
| black | AGAGGTGAGGCGGTTTGCACGTAAAACATATCAACAATAGAT | | | | | | |
| black | GCCATTAAAAATACGTTTAACGTCAGATGACAATAAACAACA | | | | | | |
| black | GCTATTAGTCTTTACGGGAGAAACAATAATAAAGTACCGACA | | | | | | |
| black | TGTACCGTAACACTTTTTGTCACAATCAGGAATACCCAAAAG | | | | | | |
| black | TGAAAGCGTAAGAAAAGTTACAAAATCGGTAATTTAGGCAGA | | | | | | |
| black | AAAAGGGACATTCTCCTGAGCAAAAGAATAGGGCTTAATTGA | | | | | | |
| black | TTACCAGTATAAAGCGGTAATCGTAAAAATCGGTGCGGGCCT | | | | | | |
| black | CTAGAAAAAGCCTGTGATAATCAGAAAAGCGCCATTCGCCAT | | | | | | |
| black | AAATAAGGCGTTAAAAATATTTAAATTGCCAGCTTTCCGGCA | | | | | | |
| black | TTCTGACCTAAATTATTAAATTTTTGTTGGGGACGACGACAG | | | | | | |
| black | ACGCGAGAAAACTTACGCCATCAAAAATGGTGTAGATGGGCG | | | | | | |
| black | AAGGAAACCGAGGACGTCATAAATATTCAACTAATGCAGATA | | | | | | |
| black | CTATATGTAAATGCCTTTCATCAACATTGGGAACAAACGGCG | | | | | | |
| black | CTTTTTAAGAAAAGGTTCAGAAAACGAGGGTAGAAAGATTCA | | | | | | |
| black | AGCAAGAAACAATGTACCCTGACTATTAATCTACGTTAATAA | | | | | | |
| black | AGAGAGATAACCCAAAAGATTAAGAGGAAAGAACTGGCTCAT | | | | | | |
| black | AACACCCTGAACAATAATTCGAGCTTCATAATTTCAACTTTA | | | | | | |
| black | AACATAAAAACAGGACAGGTCAGGATTAGAGAAACACCAGAA | | | | | | |
| black | TTTAACGTCAAAAAAAGAGGTCATTTTTCGTAACAAAGCTGC | | | | | | |
| black | GCCATATTATTTATTATAATGCTGTAGCGAGTAATCTTGACA | | | | | | |
| black | AGCGTCTTTCCAGATACGGTGTCTGGAACGGTGTACAGACCA | | | | | | |
| black | CACCCAGCTACAATAATTCTGCGAACGATAAGGGAACCGAAC | | | | | | |
| black | GGGAGGTTTTGAAGTTCGCAAATGGTCACTCCATGTTACTTA | | | | | | |
| black | ATTCTAAGAACGCGGGGCGCGAGCTGAATTGTATCATCGCCT | | | | | | |
| black | CGCCCAATAGCAAGTAGCATTACATCCACGGGGAGAGGCGGT | | | | | | |
| black | ACAAGCAAGCCGTTAGCAAAATTAAGCAGAAACCTGTCGTGC | | | | | | |
| black | TTCCAAGAACGGGTGGTTGTACCAAAAATCACATTAATTGCG | | | | | | |
| black | GCATGTAGAAACCAAGAAGCCTTTATTTGCATAAAGTGTAAA | | | | | | |
| black | AACTGGCATGATTATAGTAAAATGTTTAAGTAAGAGCAACAC | | | | | | |
| black | AAGTCCTGAACAAGCTCATATATTTTAAATTGTTATCCGCTC | | | | | | |
| black | TGTTCAGCTAATGCAGATTCAAAAGGGTGCTCGAATTCGTAA | | | | | | |
| black | AAAGGTAAAGTAATATCAATATGATATTTGCATGCCTGCAGG | | | | | | |
| black | GGCATTTTCGAGCCGGAGAGGGTAGCTATTCCCAGTCACGAC | | | | | | |
| black | GAATCGCCATATTTGTCATTGCCTGAGAGGGATGTGCTGCAA | | | | | | |

| color in cadnano file | sequence | +a-ssDNA | +s-ssDNA | +1 h/h'-ssDNA | +3 h/h'-ssDNA | +7 h/h'-ssDNA | +14 h/h'-ssDNA |
|---|---|---|---|---|---|---|---|
| black | CTGAGGCTTGCAGGAGGCTTGCCCTGACGAGAGTACCTTTAA | | | | | | |
| black | AAATCGGAACCCTAGTAACGCCAGGGTTTTTTTGAGAGATCT | | | | | | |
| black | ATATGTACCCCGGTTTTAGTATCATATGAACAATTTCATTTG | | | | | | |
| black | AAGATTGTATAAGCATAAGAATAAACACTAAATCAATATATG | | | | | | |
| black | TTGTTAAAATTCGCTAATGGTTTGAAATGTCGCTATTAATTA | | | | | | |
| black | TTTAACCAATAGGATTTCAAATATATTTGCGATAGCTTAGAT | | | | | | |
| black | CTTCCTGTAGCCAGTGATGCAAATCCAATTTATCAAAATCAT | | | | | | |
| black | AAATGCTTTAAACATAAGCAGATAGCCGCGACATTCAACCGA | | | | | | |
| black | AAAAATCAGGTCTTAAATAGCAATAGCTAAATTATTCATTAA | | | | | | |
| black | GCGGATTGCATCAACAAGAATTGAGTTAGCCATTTGGGAATT | | | | | | |
| black | GGAAGCAAACTCCAGAAGCGCATTAGACATAGCAGCACCGTA | | | | | | |
| black | TTGCTCCTTTTGATTGAAAATAGCAGCCTTAGCGTCAGACTG | | | | | | |
| black | GCTTAATTGCTGAACCCAATCCAAATAAATAGCCCCCTTATT | | | | | | |
| black | ATATGCAACTAAAGGCCTAATTTGCCAGTCACCGGAACCAGA | | | | | | |
| black | AACAGTTGATTCCCTTTATCCTGAATCTCCGCCACCCTCAGA | | | | | | |
| black | GCCAGAGGGGGTAAAGACTCCTTATTACAACGCAAAGACACC | | | | | | |
| black | TATATTTTCATTTGAGGCGTTTTAGCGAACAGGAGTAGACTT | | | | | | |
| black | TCTACTAATAGTAGCAAATCAGATATAGTCCTTTGCCCGAAC | | | | | | |
| black | CAAGGCAAAGAATTTTTATTTTCATCGTTTATCATTTTGCGG | | | | | | |
| black | CATAAAGCTAAATCATTAAACCAAGTACTTATCATCATATTC | | | | | | |
| black | AATACTTTTGCGGGATCAATAATCGGCTATATAATCCTGATT | | | | | | |
| black | TAATGTGTAGGTAAAGAACGCGCCTGTTGAAATAAAGAAATT | | | | | | |
| black | ACAGTCAAATCACCTCTGTCCAGACGACGAATATACAGTAAC | | | | | | |
| black | GATAAATTAATGCCAGTAATAAGAGAATACGGATTCGCCTGA | | | | | | |
| black | CAATACTGCGGAATAACGCAATAATAACATAGAAAATTCATA | | | | | | |
| black | ACAAAGGCTATCAGAACAACGCCAACATCGCAGAGGCGAATT | | | | | | |
| black | AGAGAATCGATGAACCAACGCTCAACAGGATGATGAAACAAA | | | | | | |
| black | GAAAGGAGCGGGCGTGTTGGGAAGGGCGCTAGCATGTCAATC | | | | | | |
| black | ACAGGGCGCGTACTAGATCGCACTCCAGTAAACGTTAATATT | | | | | | |
| black | CGATTAAAGGGATTGGATAGGTCACGTTAATTCGCGTCTGGC | | | | | | |
| black | TAATTTTTTCACGTACAACATTATTACAAATGACCATAAATC | | | | | | |
| black | TGAATTTCTTAAACCCTTATGCGATTTTAGCCCGAAAGACTT | | | | | | |
| black | CGGCTACAGAGGCTCTGACCTTCATCAATCAACATGTTTTAA | | | | | | |
| black | GCACCAACCTAAAACAGACGGTCAATCAGTAGATTTAGTTTG | | | | | | |
| black | AGTACAAGGTTTTTCCAGGGTCGGAGATAAGGTGGCATCAAT | | | | | | |
| black | GGTCCACGCTGGTTCGCTTTCCAGTCGGATAAAGCCTCAGAG | | | | | | |
| black | ATAGCCCGAGATAGCATACGAGCCGGAACAACGCAAGGATAA | | | | | | |
| black | TTCTGTATGGGATTGAATTACGAGGCATGACTGGATAGCGTC | | | | | | |
| black | AAAAACCGTCTATCTCCCCGGGTACCGAGAGAAAGGCCGGAG | | | | | | |
| green | AGCGAGAGGCTTTTATAAAAACCAAAAT | yes (edge) | | | | | |
| green | TAGTTAGCGTAACGACAGACAGCCCTCA | yes (edge) | | | | | |
| green | ATACATAAAGGTGGAACGTAGAAAATAC | yes (edge) | | | | | |
| green | CAAATATCGCGTTTAGTCAGAGGGTAATTTACCATTAGCAAG | yes (center) | | | | | |
| green | ACCATTAGATACATCCTTAAATCAAGATGAGCCGCCACCAGA | yes (center) | | | | | |
| green | AAATTTTTAGAACCAAAAATAATATCCCGGGTTAGAACCTAC | yes (center) | | | | | |
| yellow | GCTGCGCGTAACCAGGAAACCAGGCAAAGCCCCAAAAACAGG | | yes | | | | |
| yellow | TGCTTTCCTCGTTAATCTGCCAGTTTGAAAATCAGCTCATTT | | yes | | | | |
| yellow | GGAGTGAGAATAGAGGAATACCACATTCATTGAATCCCCCTC | | yes | | | | |
| yellow | AGGAGCCTTTAATTCGTTGGGAAGAAAATAGTCAGAAGCAAA | | yes | | | | |
| yellow | TGACAACAACCATCGGCTTGAGATGGTTAAGCGAACCAGACC | | yes | | | | |
| yellow | ATCTAAAGTTTTGTTTTACCAGACGACGGCAAAAGAAGTTTT | | yes | | | | |
| yellow | CACCCTCAGCAGCGATTACCCAAATCAAGCGGATGGCTTAGA | | yes | | | | |
| yellow | AGTTTCCATTAAACAGAGGACAGATGAAGTTTCATTCCATAT | | yes | | | | |
| yellow | ACTCATCTTTGACCAAATCCGCGACCTGATAACCTGTTTAGC | | yes | | | | |
| yellow | GATTGCCCTTCACCAATCGGCCAACGCGATAAATCATACAGG | | yes | | | | |
| yellow | TGGTGGTTCCGAAAATGAGTGAGCTAACCATTATGACCCTGT | | yes | | | | |
| yellow | GAGTCCACTATTAAGTTTCCTGTGTGAAATGCAATGCCTGAG | | yes | | | | |
| yellow | ACCCAAATCAAGTTGCCAGTGCCAAGCTCAACCGTTCTAGCT | | yes | | | | |
| yellow | GGGAAAGCCGGCGACAGCTGGCGAAAGGGTCTGGAGCAAACA | | yes | | | | |
| red | ACGGAATAAGTTTAGAGTTTCGTCACCATTAGTAAATGAATT | | | | yes | yes | yes |
| red | TGGTTTACCAGCGCCAGGGATAGCAAGCTCAACAGTTTCAGC | | | | | | |
| red | TTGAGGGAGGGAAGCACCCTCAGAACCGGGAATTGCGAATAA | | | | | | yes |
| red | AGGTGAATTATCACCACCGTACTCAGGAAAAAAGGCTCCAAA | | | | | | |
| red | AGAGCCAGCAAAATCGTCGAGAGGGTTGGCTTGCTTTCGAGG | | | | | yes | yes |
| red | GCCGGAAACGTCACTAGGATTAGCGGGGAGTTGCGCCGACAA | | | | | | |
| red | ATCAGTAGCGACAGACATGAAAGTATTAGATATATTCGGTCG | | | | | | yes |
| red | TAGCGCGTTTTCATCAGTTAATGCCCCCTTTTGCGGGATCGT | | | | | | |
| red | AGCGTTTGCCATCTTTTTAACGGGGTCAAACGAGGGTAGCAA | | | | | yes | yes |
| red | GCCACCACCGGAACCATACATGGCTTTTCTTTTTCATGAGGA | | | | | | |
| red | ACCGCCACCCTCAGAATGGAAAGCGCAGATGCCACTACGAAG | | | | | | yes |
| red | ACCACCACCAGAGCGGCCTTGATATTCAGAATACACTAAAAC | | | | | | |
| red | TACAAACAATTCGACATTTGAGGATTTACAAGCGCGAAACAA | | | yes | yes | yes | yes |
| red | GTTATTAATTTTAAGAGCACTAACAACTGACGGGCAACAGCT | | | | | | |
| red | AACAAAGAAACCACCAGTTGAAAGGAATGAGTTGCAGCAAGC | | | | | | yes |
| red | CTGATTATCAGATGATATCAAACCCTCAAAAATCCTGTTTGA | | | | | | |
| red | GTTTGGATTATACTCAGCAAATGAAAAATATAAATCAAAAGA | | | | | yes | yes |
| red | CATATCAAAATTATCAGTATTAACACCGCCAGTTTGGAACAA | | | | | | |
| red | GCGTAGATTTTCAGCGAACGAACCACCAAACGTCAAAGGGCG | | | | | | yes |
| red | AGTACCTTTTACATATGCGCGAACTGATCTACGTGAACCATC | | | | | | |
| red | TTGCTTTGAATACCTACGTGGCACAGACTGCCGTAAAGCACT | | | | | yes | yes |
| red | ATTCATTTCAATTAGGCCAACAGAGATATTTAGAGCTTGACG | | | | | | |
| red | CATCAAGAAAACAACAGATTCACCAGTCGAAGGGAAGAAAGC | | | | | | yes |
| red | AATTACCTTTTTTACATTTTGACGCTCAAGTGTAGCGGTCAC | | | | | | |
| red | TGAGTGAATAACCTCGCCAGCCATTGCACTTAATGCGCCGCT | | | | yes | yes | yes |
| red | ATTTTCCCTTAGAATCAAACTATCGGCCGAGCACGTATAACG | | | | | | |
| red | TAAGACGCTGAGAATTCTTTGATTAGTAGCTAAACAGGAGGC | | | | | | yes |
| red | AGGTCTGAGAGACTAAAGAGTCTGTCCATACGCCAGAATCCT | | | | | | |

(2) Virtanen, P.; Gommers, R.; Oliphant, T. E.; Haberland, M.; Reddy, T.; Cournapeau, D.; Burovski, E.; Peterson, P.; Weckesser, W.; Bright, J.; van der Walt, S. J.; Brett, M.; Wilson, J.; Millman, K. J.; Mayorov, N.; Nelson, A. R. J.; Jones, E.; Kern, R.; Larson, E.; Carey, C. J.; Polat, İ.; Feng, Y.; Moore, E. W.; VanderPlas, J.; Laxalde, D.; Perktold, J.; Cimrman, R.; Henriksen, I.; Quintero, E. A.; Harris, C. R.; Archibald, A. M.; Ribeiro, A. H.; Pedregosa, F.; van Mulbregt, P.; SciPy 1.0 Contributors; Vijaykumar, A.; Bardelli, A. P.; Rothberg, A.; Hilboll, A.; Kloeckner, A.; Scopatz, A.; Lee, A.; Rokem, A.; Woods, C. N.; Fulton, C.; Masson, C.; Häggström, C.; Fitzgerald, C.; Nicholson, D. A.; Hagen, D. R.; Pasechnik, D. V.; Olivetti, E.; Martin, E.; Wieser, E.; Silva, F.; Lenders, F.; Wilhelm, F.; Young, G.; Price, G. A.; Ingold, G.-L.; Allen, G. E.; Lee, G. R.; Audren, H.; Probst, I.; Dietrich, J. P.; Silterra, J.; Webber, J. T.; Slavič, J.; Nothman, J.; Buchner, J.; Kulick, J.; Schönberger, J. L.; de Miranda Cardoso, J. V.; Reimer, J.; Harrington, J.; Rodríguez, J. L. C.; Nunez-Iglesias, J.; Kuczynski, J.; Tritz, K.; Thoma, M.; Newville, M.; Kümmerer, M.; Bolingbroke, M.; Tartre, M.; Pak, M.; Smith, N. J.; Nowaczyk, N.; Shebanov, N.; Pavlyk, O.; Brodtkorb, P. A.; Lee, P.; McGibbon, R. T.; Feldbauer, R.; Lewis, S.; Tygier, S.; Sievert, S.; Vigna, S.; Peterson, S.; More, S.; Pudlik, T.; Oshima, T.; Pingel, T. J.; Robitaille, T. P.; Spura, T.; Jones, T. R.; Cera, T.; Leslie, T.; Zito, T.; Krauss, T.; Upadhyay, U.; Halchenko, Y. O.; Vázquez-Baeza, Y. SciPy 1.0: Fundamental Algorithms for Scientific Computing in Python. *Nat. Methods* **2020**, *17* (3), 261–272. https://doi.org/10.1038/s41592-019-0686-2.

(3) Theret, D. P.; Levesque, M. J.; Sato, M.; Nerem, R. M.; Wheeler, L. T. The Application of a Homogeneous Half-Space Model in the Analysis of Endothelial Cell Micropipette Measurements. *J. Biomech. Eng.* **1988**, *110* (3), 190–199. https://doi.org/10.1115/1.3108430.
